# Inactivation of MexT in *Pseudomonas aeruginosa* destabilizes cooperation and favors the emergence of a unique quorum sensing variant

**DOI:** 10.1101/2025.09.19.677155

**Authors:** Kiana R. Bellamoroso, Maxim Kostylev, Nicole E. Smalley, E. Peter Greenberg, Ajai A. Dandekar

## Abstract

*Pseudomonas aeruginosa*, an opportunistic gram-negative pathogen, uses a cell-cell signaling system called quorum sensing to coordinate group behaviors. Quorum sensing in *P. aeruginosa* is a model to study cooperative behaviors in populations. Several studies of cooperation have been conducted using strain PAO1. Wildtype PAO1 harbors a mutation in *mexS*, which encodes a negative regulator of the transcription factor MexT, which in turn activates many genes, including the efflux pump MexEF-OprN. We hypothesized that the PAO1 *mexS* mutation might affect cooperative behaviors. When *P. aeruginosa* is passaged daily on casein as a sole carbon source, quorum sensing is required to induce synthesis of the extracellular proteases needed to acquire carbon and energy. During growth on casein, individuals with inactivating mutations in the quorum sensing regulator LasR are reproducibly enriched in the population. These LasR mutants are cheaters that benefit from cooperatively-produced proteases and have a fitness advantage over cooperators. We passaged wildtype PAO1, PAO1 with a gene-corrected version of *mexS*, or PAO1 with a null mutation of *mexT* on casein as the sole carbon and energy source. We found that correcting the *mexS* mutation resulted in unstable cooperation: bacterial cultures failed to propagate after about 15 days, whereas the wildtype propagated for the duration of our 30-day experiment. The MexS-corrected and MexT-deficient populations also reproducibly exhibited emergence of a particular quorum sensing variant, LasR-V226I. This variant activated a subset of quorum-sensing regulated genes, suggesting an evolutionary pathway to alter *P. aeruginosa* quorum sensing regulons.

**Importance:** Quorum sensing governs cooperative activities and has been used as a model for studying cooperative behaviors in bacterial populations. *Pseudomonas aeruginosa* has two interlinked acyl-homoserine lactone quorum sensing circuits, LasR-I and RhlR-I; the relationship between these circuits is influenced by a regulator called MexT. Inactivation of MexT results in enhanced activity of the quorum sensing transcription factor, RhlR, and we found that it also destabilized cooperative behaviors in this bacterium. MexT-inactive strains also reproducibly support the emergence of a specific LasR variant, V226I, of *P. aeruginosa*, offering insight into evolutionary pressures that select for mutations in quorum sensing transcription factors and, by extension, a method for bacteria to alter the cohort of genes that are quorum-controlled.

## Introduction

*Pseudomonas aeruginosa*, an opportunistic human pathogen, uses an intercellular communication system called quorum sensing (QS) to regulate gene expression in a population density-dependent manner (1). QS in this bacterium activates the expression of a suite of genes that encode the production of extracellular products, such as proteases and rhamnolipids, that have shared benefits among the group. For this reason, QS in *P. aeruginosa* has been used as a model system for the study of cooperative behaviors (2).

*P. aeruginosa* QS consists of three complete circuits which together regulate an overlapping set of many genes (3). Two of these, the Las and Rhl systems, use acyl-homoserine lactone (AHL) signals. In the Las system, the synthase LasI produces the signal *N*-3-oxo-dodecanoyl-homoserine lactone (3OC12-HSL), which binds to the transcription factor LasR. Signal-bound LasR activates dozens of genes, including *rhlR*, thus regulating the Rhl system. The Rhl QS circuit consists of the synthase RhlI, which produces *N*-butanoyl homoserine lactone (C4-HSL) that binds to the transcription factor RhlR. A third QS circuit, the *Pseudomonas* quinolone signal (PQS) system, is also regulated by LasR and consists of quinolone signals that bind to the receptor PqsR (also known as MvfR).

This arrangement in which the LasR system regulates the PQS and Rhl QS systems has led to a hierarchal model of *P. aeruginosa* QS (4): deletion or inactivation of LasR results in a functionally quorum-off state, with minimal expression and activity of RhlR or PqsR. This model was developed primarily using the laboratory-adapted strain PAO1, but it is now well-understood that other factors can influence the QS hierarchy (5) and there is strain-to-strain variance in the level of Rhl circuit activation in LasR-null backgrounds (6).

*P. aeruginosa* strain PAO1 harbors an inactivating mutation in the gene *mexS*, which results in chloramphenicol resistance, presumably reflecting exposure to chloramphenicol prior to when this strain was isolated (7). The product of *mexS* negatively regulates the activity of the transcription factor MexT; MexT in turn is an activator of many genes, including those encoding the RND efflux pump MexEF-OprN (8, 9). We, and others, have previously shown that constitutive expression of MexT is in part responsible for the QS cascade in PAO1: null mutation of *mexT* results in a relaxed hierarchy in which RhlR and PqsR exhibit increased activity in the absence of LasR (10, 11).

In some circumstances, *P. aeruginosa* QS is cooperative: growth of the population depends on QS transcription factors, and the population benefits from activation of QS. One such condition is growth using casein as a sole carbon and energy source (12). *P. aeruginosa* requires the production of QS-regulated extracellular proteases to break down casein into constituent amino acids and peptides that can be imported into the cell and catabolized. Thus, QS null mutants in monoculture cannot grow on casein. However, in populations passaged on casein, LasR mutants reproducibly emerge and are enriched (13); these mutants are “social cheaters” that avail themselves of the proteases produced by wild-type cooperators, but do not incur the metabolic cost of QS and cooperation themselves, and therefore have a fitness advantage over cooperators.

When PAO1 is passaged daily on casein, the proportion of cheaters in the population typically stabilizes when they reach roughly 20-40% of the population (12, 14). The QS hierarchy has been implicated in this equilibrium, as PAO1 mutants lacking RhlR cannot control the frequency of cheaters (15, 16), which is due in part to RhlR regulation of hydrogen cyanide synthesis. When RhlR is inactive, the proportion of cheaters will continue to increase until they cannot support themselves and the population can no longer grow for lack of a sufficient number of cooperators (15). A contrasting example has been reported in populations of strain PA14. Others found that when PA14 is grown on casein, LasR mutants arise (as they do with PAO1) and achieve frequencies of up to 99% without the population collapsing (17).

We hypothesized that one possible explanation for this difference between PAO1 and PA14 is the functional status of *mexS* and, relatedly, the expression level of *mexT*. We explored whether QS mutants emerge when PAO1 populations with MexT deleted (MexT^−^) or the *mexS* gene repaired (MexS^+^) are passaged daily on casein. We found that either variation destabilized cooperative behavior in PAO1, unlike the prior observation with PA14. We also found that a particular variant of LasR, V226I, repeatedly emerged in these genetic backgrounds. This variant has also been shown to arise in other evolution experiments (14,16). The LasR-V226I variants are partially active, and individual bacteria harboring this variant behave differently in co-culture than in monoculture. These results provide insight into the selective pressures that stabilize cooperative behaviors and might favor the evolutionary divergence of QS transcription factors.

## Results

### Restoration of mexS, or deletion of mexT, in PAO1 destabilizes cooperating populations

To better understand the disparity in cheater abundance between PAO1 and PA14, we passaged wild-type (WT), MexS-restored and MexT-null PAO1 strains, as well as PA14, in a minimal medium containing casein as the sole carbon and energy source (casein broth) for up to 30 days. We hypothesized that restoration of MexS, or deletion of the gene encoding MexT in PAO1, would result in a high proportion of cheaters in cooperating populations, as observed with strain PA14.

The PAO1 mutant strains had an obvious phenotype. WT PAO1 grew more poorly in the early days of passage than either MexS^+^ or MexT^−^ strains, both of which grew well from the initiation of the experiment. The MexS^+^ and MexT^−^ strains also produced significantly more pigment than the WT (Fig. 1), consistent with our prior report of heightened Rhl activity in these strains (18). WT PAO1

**Figure 1.**
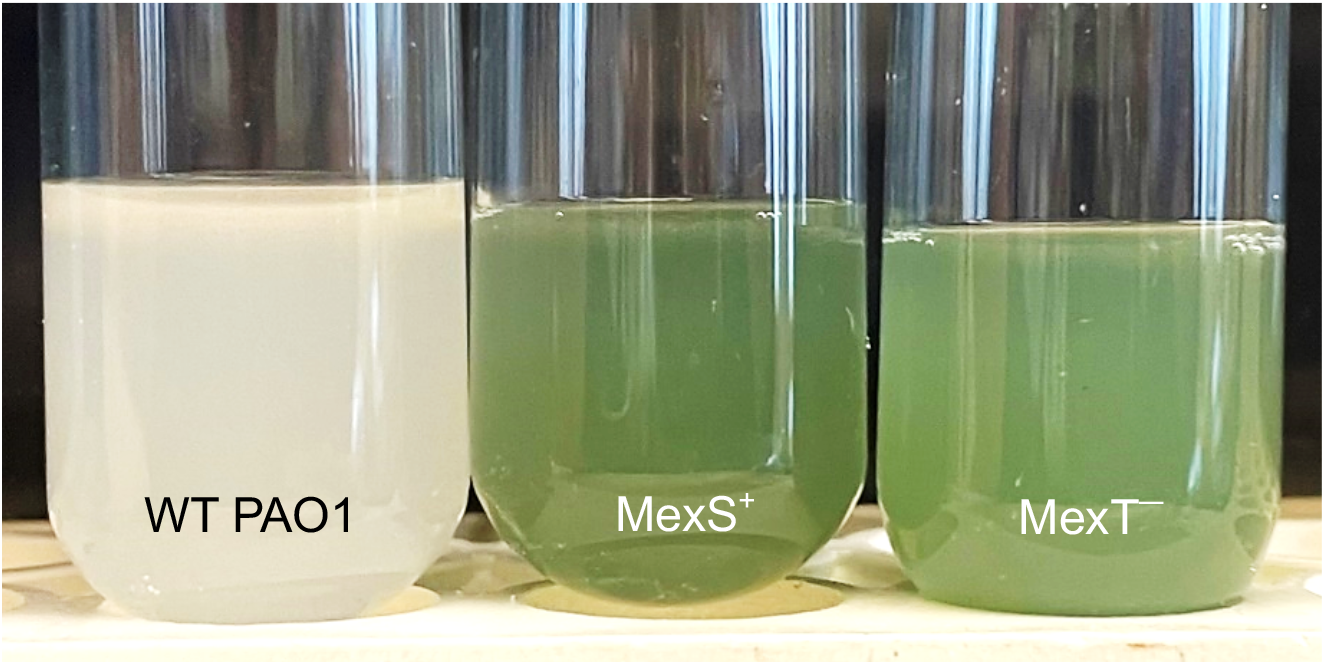
Appearance of WT PAO1, MexS^+^, and MexT^−^ after 24 hours of growth in casein broth.

We monitored the emergence of cheaters from these three backgrounds in casein broth by testing for protease production using skim-milk agar (12). As discussed above, when PAO1 is transferred daily on casein as the sole carbon and energy source, cheaters reproducibly emerge after about 10 days and often (but not always) come to an equilibrium with the WT and constitute 20-40% of the population (12, 19) (Fig. 2A). We found that restoration of MexS or deletion of MexT in PAO1 consistently resulted in cultures that were enriched for protease-negative isolates but could not be propagated after day 20 (Fig. 2B, C). There were exceptions, however: in both genetic backgrounds some populations could be passaged for all 30 days and surprisingly, protease-negative cheaters emerged and then appeared to be lost from these populations (Fig. 2B, C). In the PA14 background we observed a very high percentage of cheaters, as has been previously reported (17). However, in our experiments, the cultures could not be propagated with this burden of cheaters and all but one replicate collapsed between days 7 and 15 (Fig. 2D), likely reflecting subtle differences in the experimental conditions.

**Figure 2.**
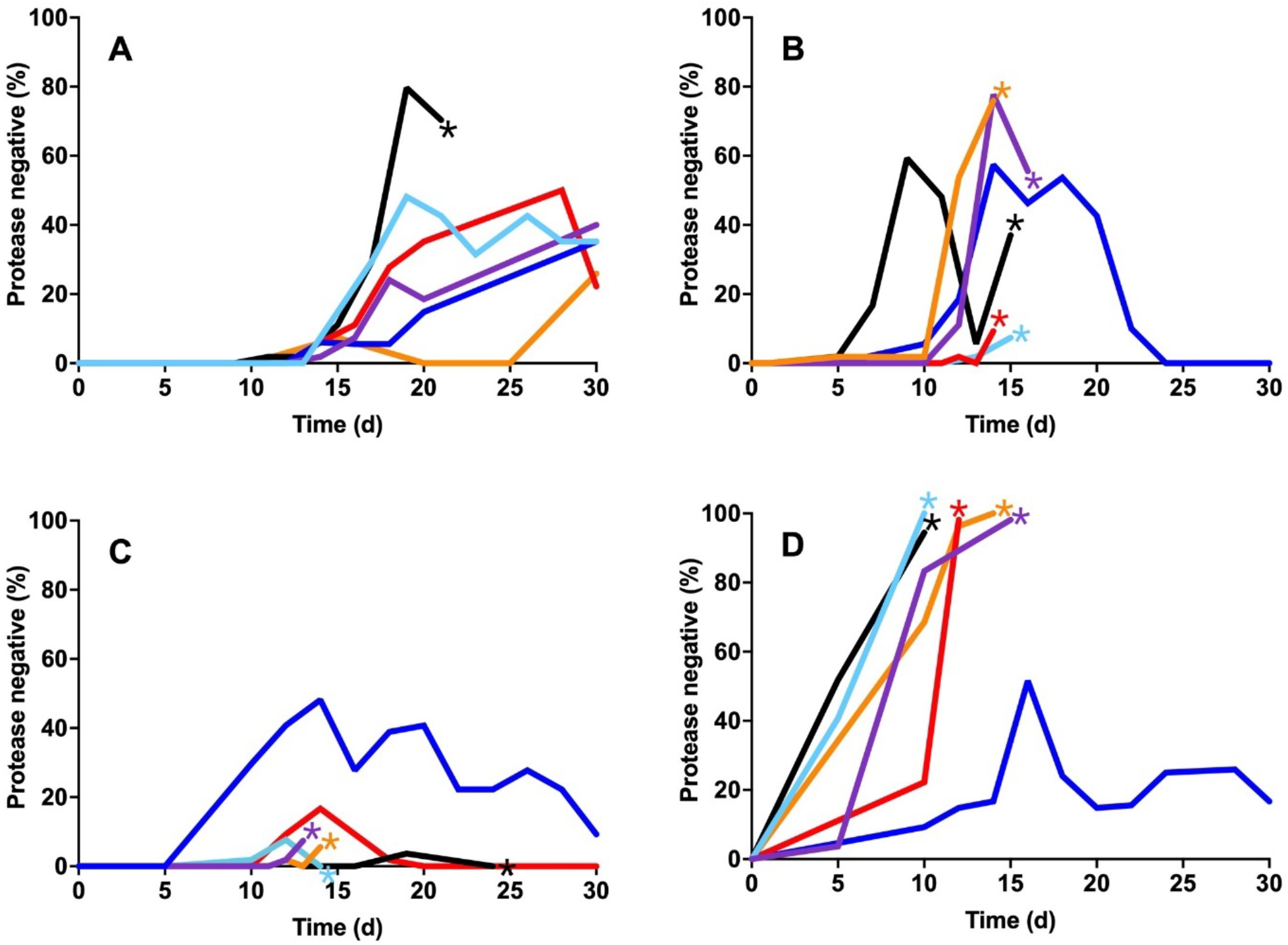
Frequency of protease-negative isolates over time in casein passage experiments. Six different isolates of PAO1 (A), MexS^+^ (B) MexT^−^ (C) and PA14 (D) were passaged daily in casein broth as described in Materials and Methods. Each line represents an individual culture. Stars denote failure of the population to propagate.

We were interested in the apparent loss of cheating in some of the MexS^+^ and MexT^−^ populations as well as the recurrent failure of other populations to propagate. We asked what mutations arose in *lasR* in the various lineages. Consistent with previous reports, the protease-negative phenotype reflected mutations in the coding sequence of *lasR* (Table S1). By day 15, an average of 26% of screened colonies in MexS-restored and MexT-knockout populations were protease-null, more than double the 12% average observed in WT strains. Most mutations in *lasR* that occur in these experiments have previously been demonstrated to be inactivating (12), and whole genome sequencing of randomly-selected protease-negative isolates from each background revealed non-synonymous *lasR* mutations (Table S3). Curiously, in this series of experiments, many protease-null isolates harbored a particular recurring mutation, G676A, encoding a V226I amino acid substitution (Table 1). After day 20, nearly 60% of isolates sequenced for *lasR* from the MexS^+^ and MexT^−^ backgrounds harbored the V226I variant. In populations where the V226I variant arose, the frequency of protease-null mutants rapidly dropped, but many of these cultures still failed to propagate (Fig. 2).

**Table 1.** *lasR* mutations found in various isolates at the final measurement timepoint.

| Background | DNA change | A.A. change | Day |
| --- | --- | --- | --- |
| MexS<sup>+</sup> | G676A | V226I | 16<sup>a</sup> |
|  | 181delC | Frameshift | 16 |
|  | G676A | V226I | 30 |
|  | G676A | V226I | 11 |
|  | G676A | V226I | 12 |
|  | G676A | V226I | 14 |
| MexT<sup>-</sup> | 89-bp deletion at C362 | del 121-150 + frameshift | 30 |
|  | G676A | V226I | 30 |
|  | G676A | V226I | 13 |
|  | G676A | V226I | 14 |
|  | G676A | V226I | 24 |

<sup>a</sup>Individual, protease-negative isolates were selected for sequencing either the day before cultures failed to propagate, or at the end of the 30-day experiment.

### MexS^+^ cooperators have a fitness disadvantage compared to both LasR-null mutants and the WT

A feature of social cheating is that cheaters have a competitive advantage over cooperators, with a diminishing benefit as their frequency increases (20). LasR mutants that arise in casein broth exhibit this negative-frequency dependent fitness advantage (12, 21). We wondered if LasR mutant cheaters would have the same competitive advantage against PAO1 MexS^+^ (or MexT^−^) as they do against WT PAO1. For simplicity we considered only the MexS^+^ strain in this and subsequent experiments, as restoration of MexS abrogates MexT activity. We asked whether the fitness advantage of LasR mutants occurred in the cocultures containing MexS^+^ cooperators and, if so, whether the advantage was due to a relative disadvantage of the cooperators or a fitness benefit to the cheaters. To answer this question, we performed competition experiments where the cooperators (LasR^+^) and cheaters (LasR^−^) did or did not have a corrected version of MexS and tested all four permutations. We observed a stronger fitness advantage for both WT Δ*lasR* and MexS^+^ Δ*lasR* mutants cocultured with a MexS^+^ cooperator than when they were with a WT cooperator (Fig. 3). Both WT Δ*lasR* and MexS^+^ Δ*lasR* showed a larger growth advantage when they were co-cultured with a MexS^+^ cooperator compared to a WT cooperator, from any starting frequency (Fig. S1).

**Figure 3.**
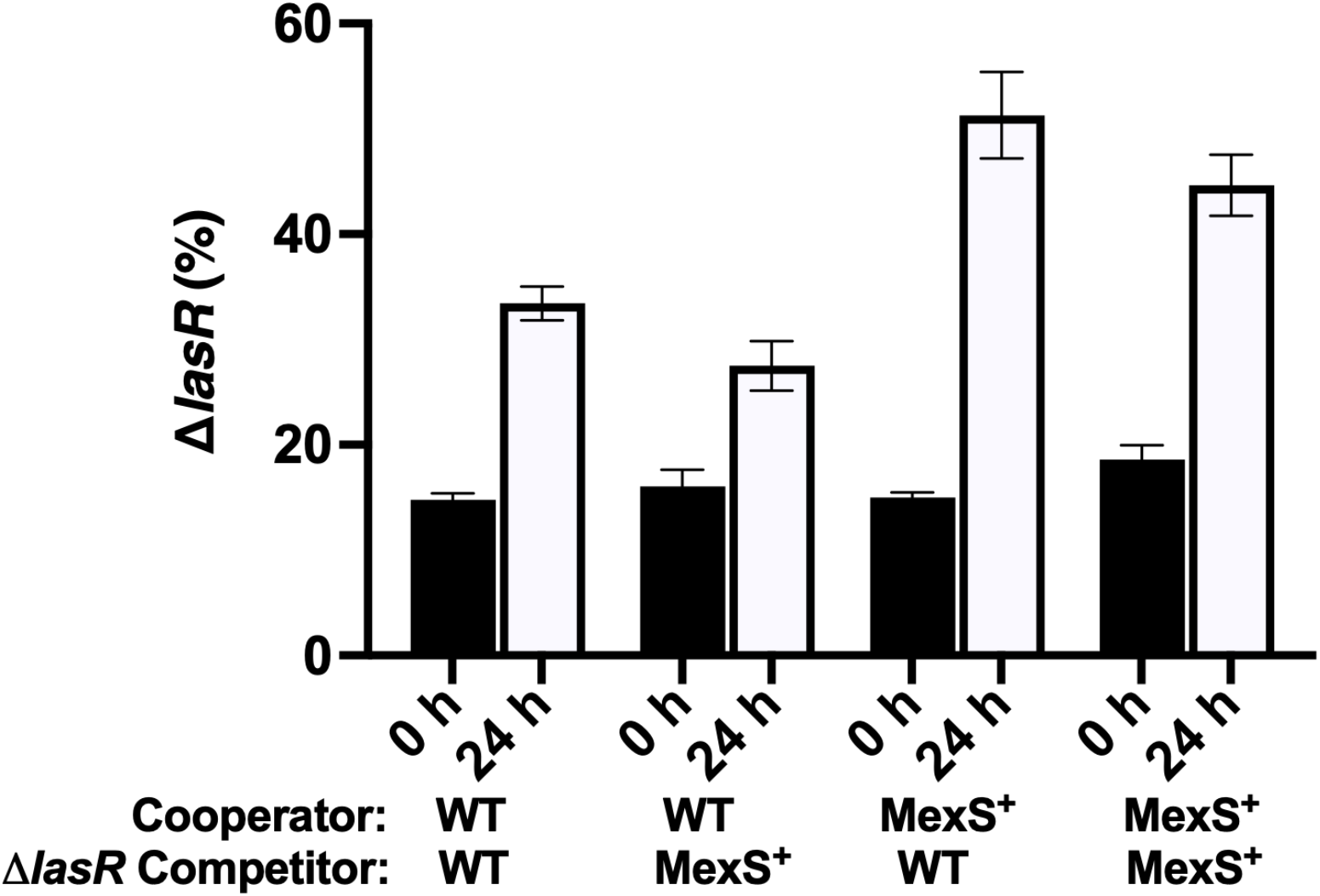
Competitions between WT and MexS^+^ cooperators and cheaters. The initial frequency of cheaters was 15%. Frequency of cheaters was enumerated at 0 h (closed bars) and 24 h (open bars) using flow cytometry. Error bars indicate SD; three trials of three replicates each.

The WT cheater also appeared to have a fitness advantage over the MexS^+^ cheater when grown with either cooperator strain: WT LasR mutants increased more than MexS^+^ LasR mutants, at all frequencies (Figs. 3 and S1), likely reflecting a fitness cost incurred by production of some QS-regulated products in LasR-mutants in the MexS^+^ background. We conclude that the *mexS* mutation present in WT PAO1 acts, unexpectedly, to stabilize cooperation in this strain.

### The V226I variant has a different phenotype than LasR-null mutants

We were intrigued by the recurrent emergence of the V226I variant in our experiments, particularly those involving the MexS^+^ and MexT^−^ mutants. The frequency at which the V226I variant emerged suggested that it had a fitness advantage over other LasR variants when grown in casein broth. We tested this hypothesis with a three-way competition experiment in which we grew WT PAO1, PAO1 Δ*lasR*, and PAO1 LasR-V226I in casein broth. We tagged PAO1 Δ*lasR* and PAO1 LasR-V226I with chromosomal copies of GFP and mCherry, respectively, and measured their relative frequency over time. We started with initial concentrations of 90% WT PAO1 and 5% of each of the LasR mutants. PAO1 LasR-V226I consistently overtook the population after 48 hours of growth. We found that on average, the LasR-V226I mutant increased from around 5% to 75% of the population within 48 hours.

We next tested whether the V226I variant behaved like a LasR-null mutant in monoculture (Fig S1). First, we grew colonies on skim-milk agar to determine if the V226I variant produced proteases, as a broad indicator of QS activity. The V226I variant is indistinguishable from PAO1Δ*lasR* in monoculture on skim milk (Fig. S2A). We also transformed PAO1Δ*lasR* or PAO1 LasR-V226I with reporter constructs in which the promoters for the quorum-sensing regulated genes *lasI, rsaL, rhlR, rhlA*, or *pqsA* were fused to *gfp* (18, 22). Gene activation in the V226I variant (as measured by GFP) is also indistinguishable from PAO1Δ*lasR* (Figs. S2B-F). We wondered if the lack of a discernable phenotype was due to lack of the QS signal 3OC12-HSL, which activates LasR. However, the addition of 3OC12-HSL had little influence on LasR-regulated gene expression, again measured using the transcriptional reporters (Fig. S3). Together, these data suggested that the V226I variant is unable to activate gene expression in monoculture.

We next asked whether the V226I mutant behaved differently than the LasR-null mutant in co-culture with the wildtype. Therefore, we co-cultured PAO1 with PAO1 LasR-V226I harboring QS transcriptional reporters (Fig. 5) and measured GFP fluorescence over time. We used the same group of reporters as in the previous experiment, and co-culture with PAO1Δ*lasR* as a control. Expression from the *lasI* and *rsaL* promoters was not substantially different between the LasR-V226I variant and the null mutant. However, there was increased transcription from the *rhlR, rhlA*, and *pqsA* promoters, consistent with activation of the Rhl and PQS QS circuits when co-cultured with WT cells. These data are consistent with the idea that some factors produced by WT cells activate expression of some QS-regulated genes in the V226I variant.

## Discussion

*Pseudomonas aeruginosa* QS is a model to study cooperative behaviors in bacterial populations. Much of the research involving QS control of cooperation with *P. aeruginosa* has involved the strain PAO1. In prior work, we and others have described how cooperation in PAO1 is stabilized by regulation of cellular factors (private goods) (14) and production of various exoproducts, including cyanide and phenazines, that “police” cheaters (15, 16, 23). These policing factors, and the means of resistance to them (16), are regulated by the QS transcriptional activator RhlR. PAO1 differs from many described strains in that it carries an inactivating mutation in *mexS* (7), ultimately resulting in constitutive MexT activity. MexT is a transcription factor that regulates multiple genes, including those encoding an efflux pump, MexEF-OprN.

Our current study demonstrates that the correction of the PAO1 *mexS* mutation or deletion of *mexT* in PAO1 led to a significant decrease in population longevity when grown in casein broth (Fig. 2). In contrast, and as previously described (12), cheaters that arose in WT populations did not usually take over the population, and the populations could be propagated for 30 days. Unlike WT PAO1, MexT^−^ and MexS^+^ populations generally failed to propagate after 10-20 days of passaging. This finding suggested that the non-functional status of the *mexS* gene in PAO1 is, unexpectedly, a stabilizing factor in maintaining QS-mediated cooperative behaviors in PAO1.

Our experiments also demonstrate that the genotype of the cooperator strain can significantly influence the fitness of LasR mutants in competition experiments. LasR mutants displayed a competitive advantage (Fig. 3) when grown in the presence of MexS^+^ cooperators compared to WT (MexS^−^) cooperators. This observation suggests that the metabolic, or possibly the QS signaling, profile of the MexS^+^ strain creates a favorable environment for cheaters. LasR mutants rapidly increased in abundance when cocultured with PAO1 MexS^+^. Together, these findings are consistent with the idea that the MexS^+^ and MexT^−^ variants incur a substantially higher metabolic cost of cooperation because they produce more QS-regulated factors (Fig. 1) (18), making it easier for cheaters to exploit them.

One occasional exception to the failure of populations to propagate that occurred in our experiments was the emergence of a particular variant of LasR, LasR-V226I. The V226I mutation is located within the DNA binding domain of LasR (24, 25). Bacteria harboring this variant, reflecting a G676A mutation in *lasR*, do not engage in QS in monoculture (Figs S2 and S3). However, the V226I variant was able to activate some QS-dependent gene expression when it was grown together with the WT (Fig. 5). The gene expression is limited to genes activated by the Rhl and PQS QS systems; the V226I variant did not activate the putatively LasR-regulated genes we studied.

In some cases (Fig 2B, C), populations in which the V226I variant arose transitioned from high frequencies of protease-null mutants to apparent cooperation over a relatively short period of time. This rapid population change is consistent with the idea that the V226I mutation provides a significant fitness advantage, while also allowing populations to potentially avoid the collapse associated with QS disruption, although we do not understand how these individuals cooperate when they form the majority of the population. It may be that the competitive advantage of the V226I variant results from its activation of known policing factors, such as hydrogen cyanide or resistance elements (15, 16). This partially functional variant occurs with low frequency in clinical and environmental isolates: a DIAMOND BLASTP analysis (26) of LasR sequences deposited in the *Pseudomonas* Genome Database (27) reveals 97 instances of the V226I variant, or a frequency of 4.9% among 239-residue (wildtype-length) LasR polypeptides. Recently, another LasR variant, A228V, has been reported to activate LasR-regulated genes, but only when its expression is enhanced by a second-site mutation (28), suggesting that variants that arise in different environments might have varied benefits, reflecting selective pressures.

The emergence of the V226I variant in our populations and its fitness advantage over both a LasR-null mutant and the WT (Fig. 4), offer insight into the evolution of acyl-homoserine lactone QS transcription factors. Prior work has focused on how new signal-receptor pairs might emerge (29, 30). Our results suggest another, mutually compatible, means by which QS circuits can change: variation in the DNA binding region of these regulators alters the suite of genes activated by them. In the case of the V226I mutant, the resultant protein does not appear to activate a LasR-regulated gene, but bacteria harboring this variant can activate RhlR- and PqsR-regulated genes (Fig. 5). Our work demonstrates that simple point mutations in the *lasR* gene, which are well-documented (6), can lead to very different QS regulatory profiles. These changes can confer a fitness benefit that results in fixation of the mutation in the population and might represent the first step on a pathway to development of a novel QS regulon.

**Figure 4.**
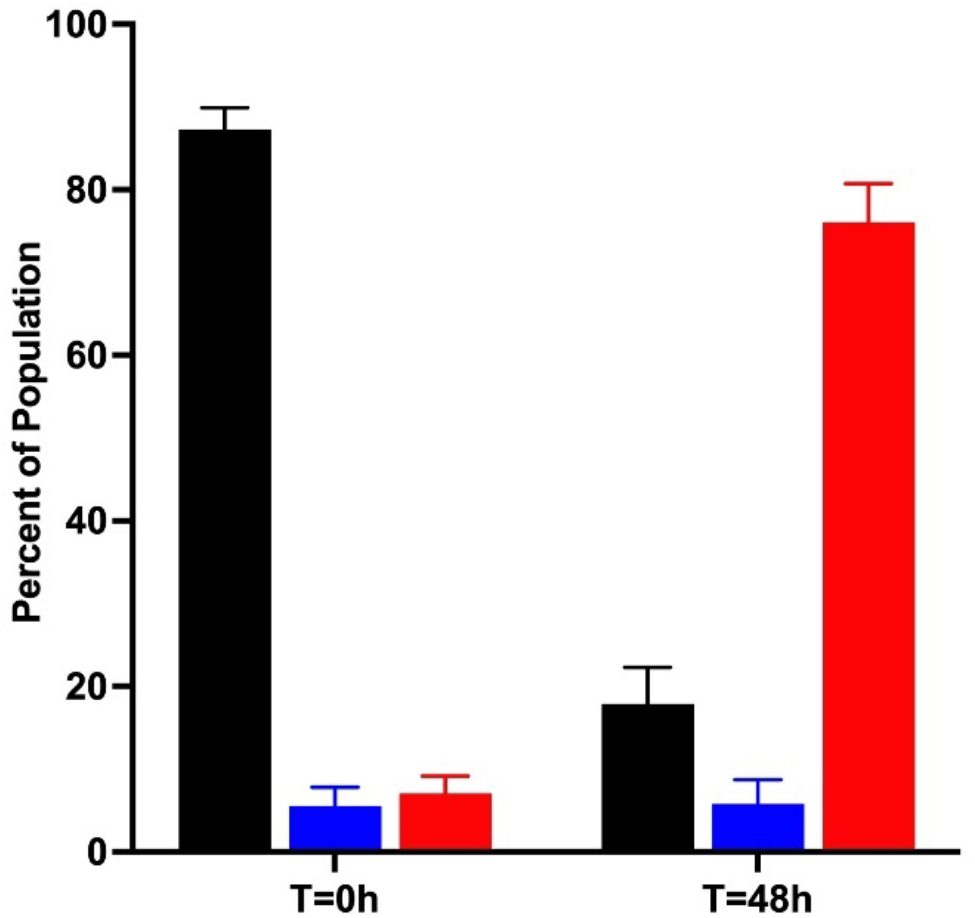
Change in population frequency of mutants and the WT after 48 h of growth in casein broth. We performed a three-way competition in casein between WT PAO1 (black bars), PAO1 Δ*lasR* (blue bars), and PAO1 LasR-V226I (red bars). Frequencies were enumerated by detection of fluorescence among individual colonies at the outset of the experiment and after 48 h of growth. Cultures were transferred to fresh media at 24 h. Data are averages of three trials of three replicates, and the error bars represent SD.

**Figure 5.**
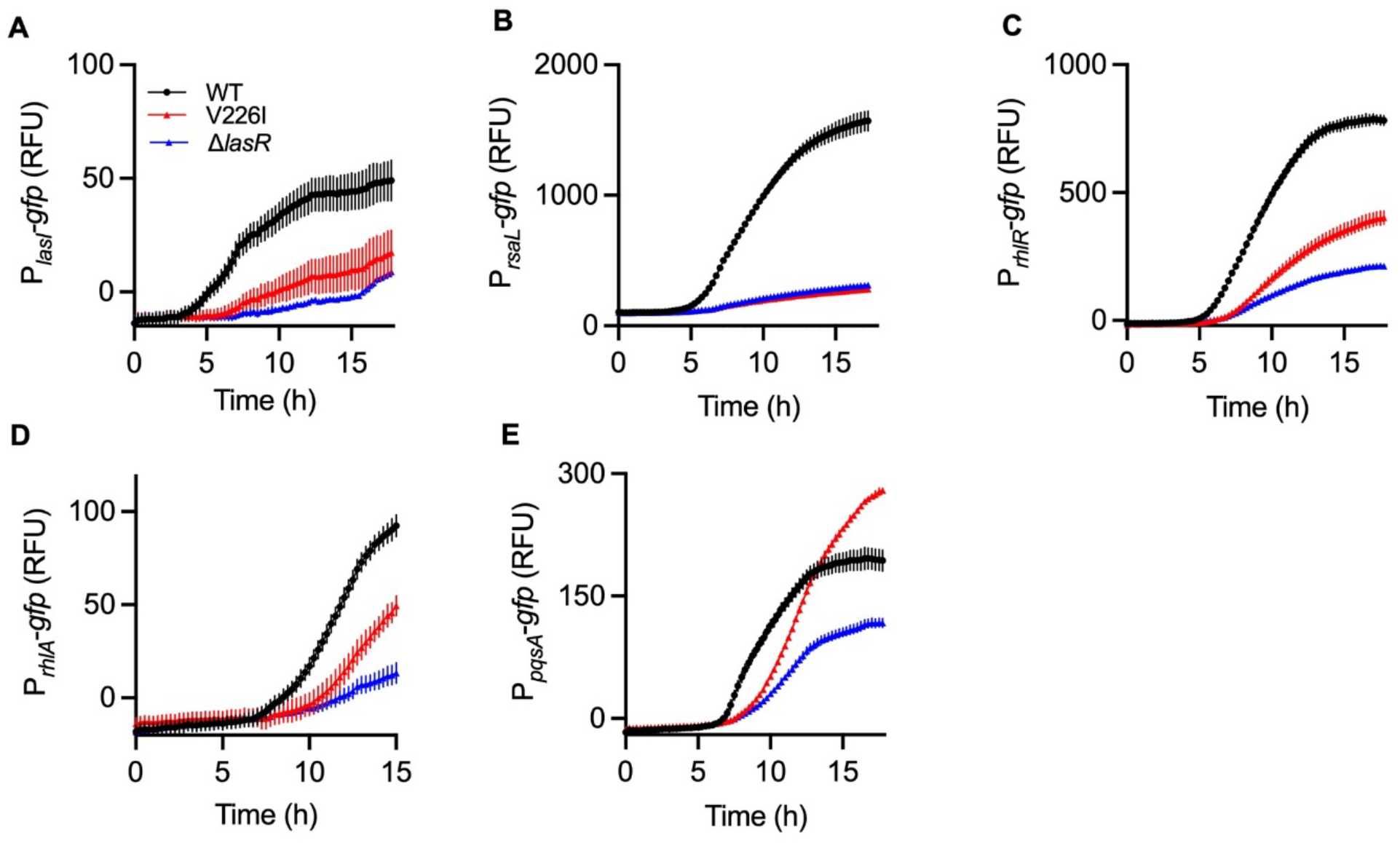
Comparison of QS-dependent reporter gene activation in LasR-V226I versus PAO1Δ*lasR*. Time course assays of gene expression of WT, LasR-V226I, and Δ*lasR* strains in co-culture with WT cells: *lasI* (**A**), *rsaL* (**B**), *rhlR* (**C**), *rhlA* (**D**), and *pqsA* (**E**). Strains containing the *gfp* reporter plasmid were inoculated with WT cells harboring an empty control plasmid at a ratio of 1:9. Data are means of four replicates, and the error bars represent SD.

## Materials and Methods

### Bacterial strains, plasmids, and media

Plasmids and strains used in these studies are listed in Supplemental Tables S2 and S3. Individual isolates were grown overnight in lysogeny broth buffered with 50 mM 3-(*N*-morpholino) propanesulfonic acid (LB) and replicates were grown in LB or a minimal medium called photosynthesis medium (31) with 1% (wt/vol) casein sodium salt (“casein broth”). When required, LB broth was supplemented with 10 ug/mL gentamicin (Gm10). Unless otherwise noted, all broth cultures were grown in 3 mL of media for 24 hours at 37°C with shaking (250 rpm). We used the homologous recombination approach in the MexS^+^ background to create our MexS^+^ Δ*lasR* strain as described (14). Briefly, DNA fragments flanking *lasR* were amplified using PCR and cloned into pEXG2. The product was used to transform *Escherichia coli* S17-1, and the pEXG2 derivatives were crossed into PAO1-derived MexS^+^. We selected transconjugants on *Pseudomonas* Isolation Agar (PIA) supplemented with 100 ug/mL gentamicin, and deletion mutants were selected using no salt LB agar containing 15% (wt/vol) sucrose. Mutant construction was confirmed by PCR.

### Casein broth passages

Three biological replicates from each strain (WT PAO1, PAO1 MexT^−^, and PAO1 MexS^+^) were grown overnight in LB. After 24 hours, 100 µL of these cultures were transferred into casein broth. We transferred 100 µl for the first 3 days, then reduced to 30 µl transfer each day for remainder of the experiment. We isolated individual colonies from the populations every two days by spreading onto LB-agar plates. From each of these spread plates, 54 individual colonies were patched onto skim milk agar plates (12) and grown overnight to ascertain protease production.

### Genotyping

Two colonies from skim milk agar plates with various protease production levels compared to WT PAO1, as evidenced by differential zones of clearing around the center of the colonies, were selected from each milk patch plate and grown overnight in LB broth. Cells were boiled and amplified for *lasR* using OneTaq polymerase buffered for GC-rich sequences. Our primers covered 157 base-pairs upstream to 140 base-pairs downstream of *lasR* (1558014 – 1559030). PCR products were purified and Sanger sequencing was performed by Azenta (formerly Genewiz). To assess for secondary mutations, we also chose two day 20+ isolates from each background and prepared genomic DNA for whole genome sequencing and variant analysis. gDNA sequencing using Oxford Nanopore technology was performed by Plasmidsaurus. Breseq (32) was used for reads mapping, alignment and variant analysis using PAO1 reference sequence GCF_000006765.1, and subsequent analysis using a University of Washington lab strain PAO1 reference to account for SNPs present in our parent strain. A cutoff of 95% variant reads with a minimum of 10 reads per locus was applied to support mutation calls.

### Competition assays

We conducted competitions between untagged MexS^+^ or MexT^−^ and a chromosomally *gfp*-tagged Δ*lasR* in casein broth. Chromosomal introduction of constitutive *gfp* and mCherry labels into the Δ*lasR* strains using mini-Tn7 was performed as described in (33). To start two-way competitions, three biological replicates of each strain were grown overnight in LB and normalized to OD_600_ 1.0. Unlabeled cooperators were combined with the Δ*lasR* strains at a starting frequency of 60%, 45%, 20%, 15%, or 10% LasR mutants to a final OD_600_ of 0.1 in 1% casein broth. The initial ratio was confirmed by flow cytometry. After 24 h of growth the frequency of Δ*lasR*-*gfp* or MexS^+^ Δ*lasR*-*gfp* in the populations was measured by flow cytometry using an Accuri C6, as previously described (34). For our three-way competition, we grew unlabeled WT, chromosomally *gfp*-tagged Δ*lasR*, and mCherry-tagged V226I overnight in LB and then combined in a ratio of 90% WT to 5% of each of the cheaters to a final OD_600_ of 0.1 in 1% casein broth. 100 µl of culture was transferred after 24 hours and dilution plates were made at time 0 and after 48 h. Colonies from these plates were then patched into a 96-well clear flat-bottom plate with 200 µl LB per well and incubated with shaking at 37°C overnight. We then measured the OD_600_ and fluorescence in a Biotek Synergy H1 plate reader (mCherry excitation 587, emission 620, and GFP excitation 485 nm, emission 528 nm).

### Gene expression in mono- and co-cultures

For both sets of gene expression assays, *P. aeruginosa* strains were electrotransformed with the transcriptional reporter plasmids P_*lasI*_*-gfp*, P_*rsaL*_*-gfp*, P_*rhlR*_*-gfp*, P_*rhlA*_*-gfp* and P_*pqsA*_*-gfp* or a promoter-less *gfp* reporter plasmid P_MCS_-*gfp* to control for background fluorescence (Table S1). For each strain, single colonies were inoculated in 3 mL LB-MOPS Gm10 and grown overnight. Following incubation, 30 µl of overnight culture were transferred into 3 mL of fresh LB-MOPS Gm10 and incubated for 2-3 doublings, or to an OD_600_ of 0.1–0.3. Monocultures were then diluted 1:10 in PBS and dispensed into duplicate wells of a 48-well clear flat-bottom plate to a final volume of 300 µl per well. In experiments with added signal, *N*-3-oxo-dodecanoyl-homoserine lactone (Cayman Chemical) in acidified ethyl acetate was added to a final concentration of 5 µM. For measurements of gene expression in co-culture experiments, WT cells carrying P_MCS_-*gfp* (blank) were combined with a strain of interest carrying either a transcriptional reporter or blank plasmid at a ratio of 9:1. Plates were incubated with double orbital shaking at 37°C in either a Biotek Synergy H1 plate reader or a Tecan Spark microplate reader with OD_600_ and GFP fluorescence (excitation 485 nm, emission 528 nm) measured every 15 minutes for 16 h. The average fluorescence values from co-cultures containing strains carrying only blank plasmids were subtracted from values obtained in co-cultures containing reporter plasmids.

### V226I variant frequency analysis

We searched for occurance of the LasR V226I protein in sequences deposited in the *Pseudomonas* Genome Database (pseudomonas.com) (27). Using the PAO1 LasR amino acid sequence as a reference, we performed a DIAMOND BLASTP (26) analysis of all *P. aeruginosa* genomes, including partial assemblies. With an E-value cutoff of 1e^−32^, default sensitivity, alignment query coverage cutoff of 30% and sequence identity cutoff of 70% we found 6,736 LasR sequences, 4,119 of which were identical to PAO1. Of the remaining 2,617 sequences, 1,963 encoded for 239-residue proteins, which is the same length as PAO1 LasR. A multiple sequence alignment of those 1,963 sequences was queried for the V226I mutation using the amino acid sequence “SRRIAA” (residues 223-228), yielding 97 polypeptides.

## Acknowledgements

This work was supported by NIH grants R01 AI177575 and R35 GM152107 (to AAD), and R35 GM136218 (to EPG). MK was supported by grants from the Cystic Fibrosis Foundation (KOSTYL20G0) and Gilead Sciences (A161091).

## Notes

### Competing Interest Statement

The authors have declared no competing interest.

