## Supplemental Figure 1 for "Inactivation of MexT in *Pseudomonas aeruginosa* destabilizes cooperation and favors the emergence of a unique quorum sensing variant"

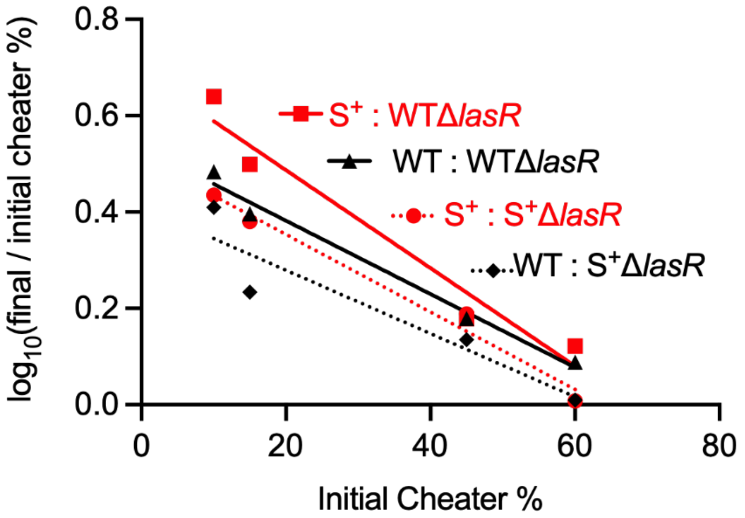


**Figure S1:** Competitive indices from coculture competitions with varying starting frequencies. WT (black) or MexS$^+$ (red) cooperators were competed against LasR-null mutants without (solid lines) or with (dotted lines) the MexS$^+$ allele. Competitions started with the initial concentration of cheaters at 10%, 15%, 45%, and 60% of the population, and the data points represent frequencies at 24 h.
