## Supplemental Figure 2 for "Inactivation of MexT in *Pseudomonas aeruginosa* destabilizes cooperation and favors the emergence of a unique quorum sensing variant"

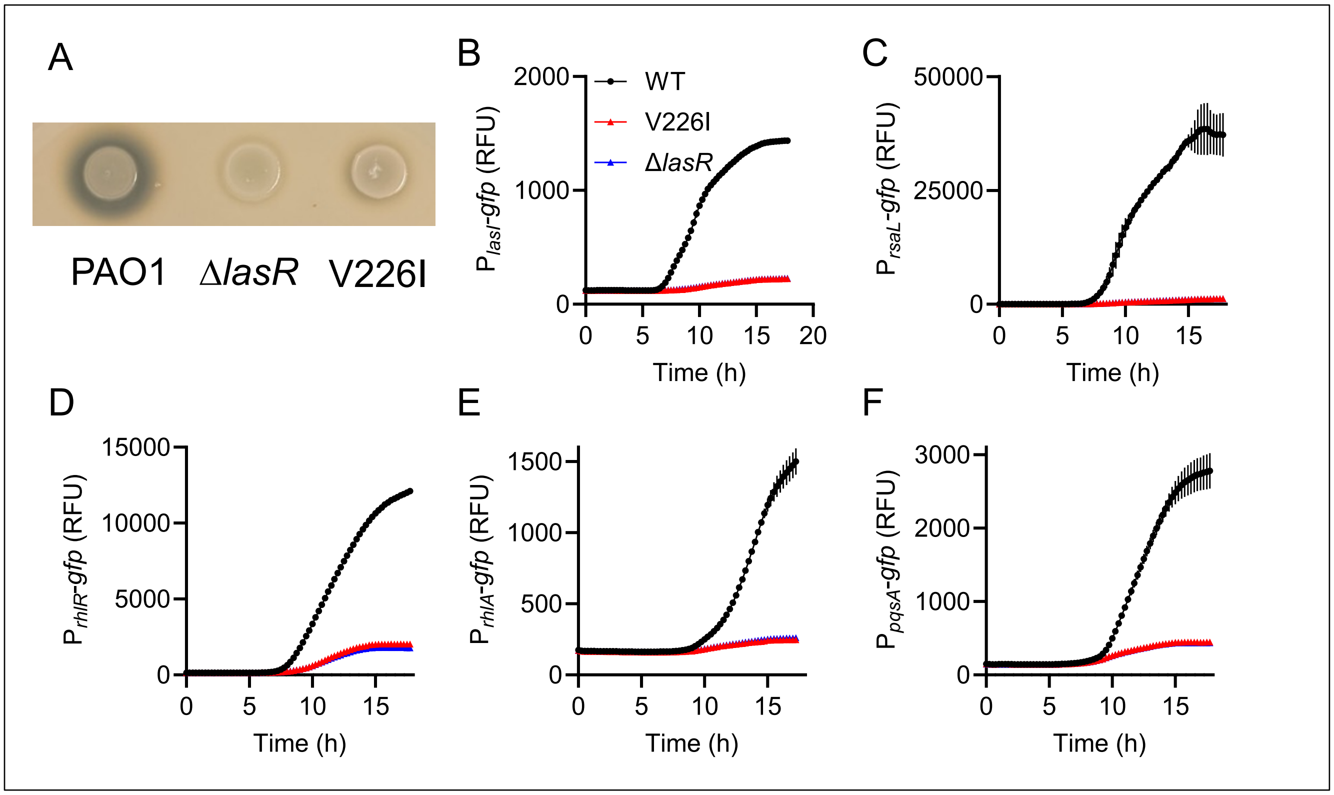


**Figure S2.** WT, LasR-V226I, and Δ*lasR* strain phenotypes in pure cultures. **A**) Colonies on skim milk agar. The zones of clearance represent protease production. **B-F**) Time course assays of gene expression in LB: *lasI* (**B**), *rsaL* (**C**), *rhlR* (**D**), *rhlA* (**E**), and *pqsA* (**F**). Where blue curves are not visible, they underlie the red curves. Data are means of four replicates, and the error bars represent SD.
