## Supplemental Figure 3 for "Inactivation of MexT in *Pseudomonas aeruginosa* destabilizes cooperation and favors the emergence of a unique quorum sensing variant"

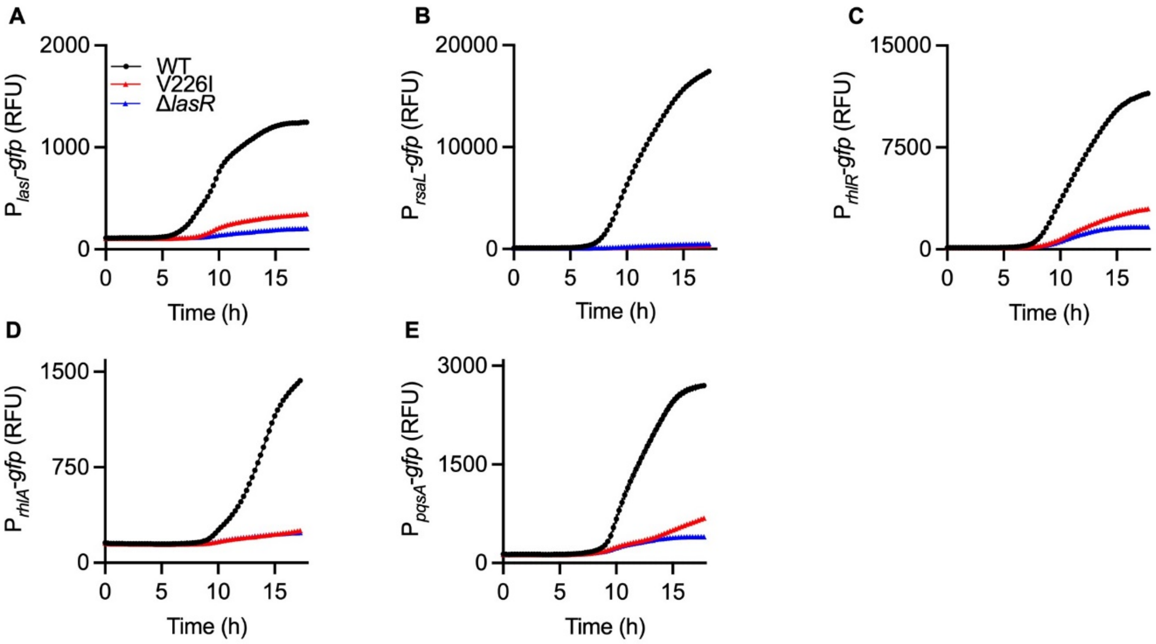


**Figure S3.** Time course assays of gene expression in LB supplemented with 5 µM of 3OC12-HSL: *lasI* (**A**), *rsaL* (**B**), *rhlR* (**C**), *rhlA* (**D**), and *pqsA* (**E**). Data are means of four replicates. Error bars representing SD are too small to see.
