## Supplemental Table 1 for "Inactivation of MexT in *Pseudomonas aeruginosa* destabilizes cooperation and favors the emergence of a unique quorum sensing variant"

**Table S1**. Whole genome sequencing of isolates from each background to identify mutations in genes other than *lasR.*

| Background[a] | Day of collection | Protease production[b] | LasR sequence[c] | Other mutations[d] | DNA change[e] | AA change[f] |
|---|---|---|---|---|---|---|
| WT | 28 | Negative | Wildtype | *gacA* | C350T | Q117*[g] |
| | | | | *psdR* | C31T | R11C |
| | | | | *pilB* | T665A | L222H |
| | | | | *rpoA* | A902G | E301G |
| WT | 28 | Negative | V226I | *psdR* | G73C | A25P |
| MexS+ | 30 | Positive | V226I | *gacA* | ∆13 bp | Frameshift, V7D |
| | | | | *psdR* | A490G | N164D |
| | | | | *pilQ* | C1990T | Q664* |
| MexS+ | 20 | Negative | F210L | *fleQ* | ∆12bp | Frameshift, G226S |
| MexT— | 30 | Positive | V226I | *gacS* | ∆1 bp | Frameshift, N267T |
| | | | | *psdR* | A479T | H160L |
| | | | | *pilQ* | G1161A | Q329* |
| | | | | *pvcC* | C854T | A285V |
| MexT— | 30 | Negative | L160* | *fleQ* | C527T | S176F |
| | | | | *psdR* | G398A | G133E |

[a] Background of isolates at initiation of casein passaging experiment.
[b] Protease production as determined by patching single colonies onto skim milk and incubating overnight at 37˚C. Positive colonies generated a zone of clearing around the colony, indicating digestion of milk protein by proteases, while negative colonies did not.
[c] Protein sequence change from wildtype PAO1 LasR (1).
[d] Single nucleotide polymorphisms (SNPs) and insertion/deletion mutations identified by *breseq* variant analysis (2) with a read frequency of 95% or higher using Stover PAO1 as the reference. SNPs called in the initial analysis were further compared to the UW parent PAO1 reference sequence to identify mutations present in the lab strain before starting the passaging experiment.
[e] For genes with mutations, loci of nucleotide changes with respect to CDS start as identified at pseudomonas.com (3).
[f] Amino acid change resulting from mutation in protein coding sequence.
[g] Asterisks denote an amino acid change resulting in an early stop codon at the referenced residue.

**Supplemental Refences**

10. Feng X, Kostylev M, Dandekar AA, Greenberg EP. Dynamics of cheater invasion in a cooperating population of *Pseudomonas aeruginosa*. Sci Rep. 9(1):10190. doi: 10.1038/s41598-019-46651-5.

11. Kostylev M, Kim DY, Smalley NE, Salukhe I, Greenberg EP, Dandekar AA. Evolution of the *Pseudomonas aeruginosa* quorum-sensing hierarchy. Proc Natl Acad Sci U S A. 116(14):7027-7032. doi: 10.1073/pnas.1819796116.

12. Simon R, Priefer U, Pühler A. 1983. A broad host range mobilization system for in vivo genetic engineering: transposon mutagenesis in gram negative bacteria. Bio/Technology 1:784–791.
