## Supplemental Table 2 for "Inactivation of MexT in *Pseudomonas aeruginosa* destabilizes cooperation and favors the emergence of a unique quorum sensing variant"

**Table S2.** Plasmids used in this study.

| Plasmid | Description | Source |
|---|---|---|
| pBBR1MCS-5 | Broad-host-range expression plasmid, $Gm^R$ | (4) |
| pBBR-MCS-*gfp* | Promoter-less *gfp* transcriptional reporter, $Gm^R$ | (5) |
| $pP_{lasI}$-*gfp* | pBBR1MCS-5 with *lasI* promoter fused to *gfp*, $Gm^R$; encodes -282 to +223 relative to the start of *lasI* and includes the complete *rsaL* binding site | (6) |
| $pP_{pqsA}$-*gfp* | pBBR1MCS-5 with *pqsA* promoter fused to *gfp*, $Gm^R$; encodes -429 to +3 relative to the start of *pqsA* | (5) |
| $pP_{rhlA}$-*gfp* | pBBR1MCS-5 with *rhlA* promoter fused to *gfp*, $Gm^R$; encodes -500 to +31 relative to the start of *rhlA* | (6) |
| $pP_{lasB}$-*gfp* | pBBR1MCS-5 with *lasB* promoter fused to *gfp*, $Gm^R$ | (6) |
| pUC18T-mini-Tn7T-Gm-*gfp* | Mini-Tn7-based vector for chromosomal integration of *gfp* and gentamicin resistancecassette at neutral *att* site; $Gm^R$ | (7) |
| pTNS2 | Helper plasmid for mini-Tn7-based chromosomal integration | (7) |
| pUC18T-mini-Tn7T-Gm-mCherry | Mini-Tn7-based vector for chromosomal integration of mCherry and gentamicin resistance cassette at neutral *att* site; $Gm^R$ | (8) |
| pEXG2-Δ*lasR* | pEXG2 allelic exchange vector with sequence for 711 bp deletion of *lasR*; preserves last 6 bp of *rsaL*; *sacB,* $Gm^R$ | (9) |
| pEXG2-LasR-V226I | pEXG2 allelic exchange vector to introduce the G676A mutation in *lasR*; *sacB,* $Gm^R$ | This work |

$Gm^R$: Gentamicin resistance
