## Supplemental Table 3 for "Inactivation of MexT in *Pseudomonas aeruginosa* destabilizes cooperation and favors the emergence of a unique quorum sensing variant"

**Table S3.** Strains used in this study.

| Strain | Description | Source |
|---|---|---|
| ***P. aeruginosa*** | | |
| PAO1 | Wild type laboratory strain with A745G mutation in *mexS* resulting in constitutive MexT activity | (1) |
| PAO1 Δ*lasR* | PAO1 derivative with unmarked, in-frame *lasR* deletion | (9) |
| PAO1 MexT— | PAO1 derivative with unmarked, in-frame *mexT* deletion | (6) |
| PAO1 MexS+ | PAO1 derivative with restored *mexS* (A745) | (6) |
| PAO1 MexS+ Δ*lasR* | PAO1 MexS+ with unmarked, in-frame *lasR* deletion | This work |
| PAO1 LasR-V226I | PAO1 derivative with partially functional LasR variant V226I | This work |
| PAO1 ΔlasR *gfp* | PAO1 Δ*lasR* with chromosomally integrated, constitutively expressed *gfp* | (10) |
| PAO1 Δ*lasR* mCherry | PAO1 Δ*lasR* with chromosomally integrated, constitutively expressed mCherry | (11) |
| PAO1 MexS+ Δ*lasR* *gfp* | PAO1 MexS+ Δ*lasR* with chromosomally integrated, constitutively expressed *gfp* | This work |
| PAO1 LasR-V226I mCherry | PAO1 LasR-V226I with chromosomally integrated, constitutively expressed mCherry | This work |
| ***E. coli*** | | |
| NEB5α | *fhuA2 Δ(argF-lacZ)U169 phoA glnV44 Φ80 Δ(lacZ)M15 gyrA96 recA1 relA1 endA1 thi-1 hsdR17* | NEB |
| S17-1 | *recA* pro *hsdR* RP4-2Tc::Mu-Km::Tn7 | (12) |
